# CAR-T Engager Proteins Optimize Anti-CD19 CAR-T Cell Therapies for Lymphoma

**DOI:** 10.1101/2022.05.30.494010

**Authors:** Lihe Su, Lan Wu, Roy R. Lobb, Paul D. Rennert, Christine Ambrose

**Author notes:** Corresponding Author: Paul D, Rennert, Aleta Biotherapeutics, 27 Strathmore Rd, Natick MA 01746 USA; 1-508-282-6370; ORCID ID 0000-0002-4573-3764; on. Twitter @PDRennert. **Declaration of interest:** All authors are employess and own stock and/or stock options in Aleta Biotherapeutics. In accordance with Journal and Publisher guidelines the authors declare this financial interest and further declare that research conducted may benefit Aleta Biotherapeutics. We have adopted ethical guidelines to prevent conflicts of interest from influencing our research. **Author Contributions:** Lihe Su and Lan Wu designed experiments and generated experimental date. Roy R. Lobb and Paul D. Rennert conceived of the experimental approach and application to specific indications; Paul D Rennert authored the manuscript; Roy R Lobb and Christine Ambrose edited the manuscript. Christine Ambrose designed and created specific therapeutics and oversaw research. The order of authorship for the co-first authors was determined by last name in alphabetical order. These authors contributed equally to the work described (Lihe Su, Lan Wu).

## Abstract

B cell lymphoma therapy has been transformed by CD19-targeting cellular therapeutics that induce high clinical response rates and impressive remissions in relapsed and refractory patients. However, approximately half of all patients who respond to CD19-directed cell therapy relapse, the majority within six months. One characteristic of relapse is loss or reduction of CD19 expression on malignant B cells. We designed a unique therapeutic to prevent and reverse relapses due to lost or reduced CD19 expression. This novel biologic, a CAR T Engager, binds CD20 and displays the CD19 extracellular domain. This approach increases the apparent CD19 antigen density on CD19-positive/CD20-positive lymphoma cells, and prevents CD19 antigen-oss induced relapse, as CD19 bound to CD20 remains present on the cell surface. We demonstrate that this novel therapeutic prevents and reverses lymphoma relapse *in vitro* and prevents CD19-negative lymphoma growth and relapse *in vivo*.

## Introduction

Chimeric Antigen Receptor (CAR) transduced autologous T cells targeting the B cell malignancy antigen CD19 illustrate the great promise of cell therapy. Anti-CD19 CAR-T cells (CAR-19) induced high response rates and durable remissions in relapsed and refractory chronic and acute lymphocytic leukemia (CLL and ALL) and non-Hodgkin lymphoma (NHL) patients (1). Long-term follow-up studies have demonstrated curative potential, with some patients reaching 10 years cancer-free (2–6).

CAR-19 effectiveness is tempered by several critical issues. The cost of therapy remains high, and specialized infrastructure can limit where CAR therapies can be created or administered (7). Preparative apheresis, consolidation, lymphodepletion and toxicities limit the use of CAR-T therapy in frail patients. Side effects from CAR-19 treatment include grade 3+ cytokine release syndrome and immune effector cell-associated neurotoxicity syndrome that can lead to prolonged hospitalization and additional cost of care (8). More than half of responding patients subsequently relapse, often within a few months of treatment (9, 10). Many patients relapse due to loss or downregulation of the CD19 protein on their leukemia or lymphoma cells (11).

Antigen escape mechanisms are common across therapeutic modalities in cancer treatment (12, 13). CAR-19 T cells put intense natural selection pressure on the malignant B cells, driving emergence of clones with lost or reduced CD19 expression. Antigen loss can occur via mutational events that prevent expression of all or part of the CD19 extracellular domain (ECD), and via transcriptional downregulation of expression (14, 15). Other mechanisms of escape from CAR-19 therapies include lineage switch to myeloid cell leukemia, and trogocytosis, a process by which CD19 is stripped from target cells by the CAR (16, 17).

We previously presented a class of molecules called bridging proteins (18, 19) or CAR-T Engagers (CTE) that can be expressed by cells transduced with viral vectors, or can be expressed in mammalian cell culture *in vitro* and purified to create biologics for injection. Here we describe a CTE biologic that increases CD19 antigen density on target lymphoma cells regardless of their level of CD19 expression. This biologic contains three functional domains: a modified CD19 ECD, an anti-CD20 binding domain and an anti-albumin binding domain. The protein, termed a CAR-19-CD20 T cell Engager (CTE-19.20), binds to CD20 and displays the CD19 ECD. CTE-19.20 proteins potently triggered CD19-negative lymphoma cell death in the presence of CAR-19 T cells *in vitro* and prevented antigen-loss relapse in an *in vitro* model of lymphoma escape from CAR-19 therapy. Using *in vivo* modeling we show that CTE-19.20 protein given alongside CAR-19 T cells prevented CD19-negative lymphoma expansion, eliminated disease, and significantly impacted survival. CTE-19.20 was readily expressed and secreted by transfected mammalian cells, was efficiently purified and demonstrated favorable biophysical properties. This therapeutic is in development to treat patients receiving CAR-19 therapeutics for B cell lymphoma and leukemia.

## Methods

### Production of anti-CD19 CAR-T cells

Anti-CD19 CAR-T cells were produced and characterized as previously described (18), except that some preparations were made using donor T cells that were activated for 48 hours before virus infection using LentiBoost (5 μl each of solutions A and B per ml of cells transfected; Fisher Scientific). 100 units/ml IL-2 was added at the start of the activation procedure and with every media change during the expansion phase.

### Mammalian cell culture

The JeKo-1 mantle cell lymphoma (MCL) and Ramos Hodgkin lymphoma cell lines were obtained from ATCC, which verifies cell line identity using short tandem repeat markers. Cells were cultured in complete media (RPMI plus 10% FBS, VWR). The JeKo-1 cell line expressing luciferase was generated using HLUC-Lv105 lentiviral particles (GeneCopoeia) and selected with puromycin. The cell lines were tested for mycoplasma (Invivogen) and were negative.

To knockout (KO) CD19 gene expression we used CRISPR/Cas9 technology. Four different CRISPR guide RNAs and Cas9 (Synthego) were combined at a 1:9 ratio and then added to luciferase-stable JeKo-1 cells suspended in Buffer R (Neon Transfection System, Thermo Fisher). The suspension was pulse electroporated at 1550 volts for 30 milliseconds, then the cells were plated into complete media to recover for 3 days. CD19 KO efficiency was determined by flow cytometry. A CD19 exon 2 guide RNA produced the highest number of KO cells and was plated using limiting dilution for single clone isolation. Expanded clones were re-assayed for CD19 expression. One clone, 1C8, was used for all studies.

### Construction and purification of CTE19.20

Previously we described a bridging protein containing a stabilized CD19 ECD linked to an anti-CD20 scFv derived from the anti-CD19 monoclonal antibody (MAb) Leu16 (20). Here we used three domains: an anti-CD20 VHH derived from a llama single domain antibody, the stabilized CD19 ECD called NT.1 (20), and the anti-albumin llama VHH, Alb8 (21). The construct, CTE-19.20, was cloned into mammalian cell expression vectors and transiently transfected into Expi293 cells (1 liter scale, Bon Opus Biosciences) or CHO cells (3 liter scale, Wuxi Biologics). The construct expressed in Expi293 cells contained a 6xHis-tag at the C-terminus (CTE-19.20-His).

Secreted CTE-19.20-His protein was purified by His-NTA column (Shanghai YuanYe) chromatography followed by monomer isolation by size exclusion chromatography (SEC, Superdex 200, GE) on an AKTA explorer (GE) by Bon Opus Biosciences. Research grade, CHO-derived protein (CTE-19.20-RG), was purified at WuXi Biologics using protein A-column and CEX chromatography, achieving 95.0% purity by SEC-HPLC. Purified proteins were also analyzed by SDS PAGE under reducing and non-reducing conditions using a 4-12% Bis-Tris Gel (NuPage). The endotoxin level was <1EU/mg.

### Binding and cytotoxicity assays

ELISA assays were developed to characterize binding to the targeted proteins. We used His-tag based ELISAs as previously described (20), and also developed assays that allowed simultaneous detection of multiple binding events. ELISA plates (Thermo Fisher) were coated overnight at 4°C with 1 μg/ml anti-CD19 MAb FMC63 (Novus) and blocked with 200 μl/well 0.3% non-fat milk in Tris buffered saline (TBS) for one hour at room temperature. CTE-19.20 proteins were added in 1% BSA in TBS, in decreasing concentrations, and incubated for 1 hour. The plate was washed 3 times with TBS. Either 0.5 μg/ml biotinylated human albumin or 0.5 μg/ml biotinylated CD20-“nanodisc” (both from Acro Biosystems) was then added in 1% BSA in TBS for 1 hour, the plates were washed again, followed by incubation with streptavidin-HRP and TMB peroxidase substrate (Thermo Fisher) to detect binding.

Cell staining was performed at 4°C. CTE-19.20 proteins, diluted in FACS buffer (PBS + 1% BSA + 0.1% sodium azide), were incubated with cells for 30 minutes, washed, then incubated with PE-coupled anti-CD19 MAb FMC63 (EMD Millipore) (5 μl/sample) for 30 minutes, washed again, then fixed with a final concentration of 1% paraformaldehyde (Thermo Scientific). Samples were read on a Accuri C6 Plus flow cytometer (BD) and results analyzed using FlowJo. In some experiments incubations were done in up to 100% human serum in order to evaluate the effect of excess human albumin on binding to cells.

Two assays were used to assess cross reactivity to murine proteins. For flow cytometry analyses, murine and human CD20 cDNA expression constructs (GenScript) were transiently transfected into 293T cells using lipofectamine (ThermoFisher). The cells were removed using Accutase (Corning), and binding to CTE-19.20 proteins was detected using the flow cytometry assay. Second, murine albumin was coated onto an ELISA plate and increasing concentrations of CTE-19.20-His were added to the plate and incubated for 1 hour at room temperature. An anti-His-tag antibody coupled to HRP (Thermo Scientific) was used for enzymatic detection. The blocking, dilution buffer and wash buffers were as detailed above.

Cytotoxicity assays were run as previously described (18, 20) using lymphoma cells, CAR-19 T cells and CTE proteins. In some experiments the media was supplemented with 50% human serum in order to evaluate the effect of excess human albumin on cytotoxicity.

### Effect of CTE.19-20 proteins on cells

To assess whether soluble CAR-T Engager protein could influence direct CD19 cytotoxicity, JeKo-1 cells were cultured with CAR-19 T cells alone or with up to 1 μg/ml CTE-19.20-RG, a saturating amount. The E:T ratios were 3:1 and 1:1. CAR-19-mediated cytotoxicity was measured after 48 hours. Down-regulation of cell surface CD20 was measured using JeKo-19KO cells incubated at 4°C or 37°C for varying lengths of time with CTE-19.20-RG, then analyzed by flow cytometry as described above. To evaluate whether excess CTE-19.20-RG could saturate target proteins, an order of addition assay was performed using the cytotoxicity assay. Three incubation conditions were evaluated: (1) target cells, CTE-19.20-RG protein and CAR-19 T cells were added at the same time; (2) target cells and CTE-19.20-RG protein were incubated together at 37°C for 10 minutes and then added to CAR-19 T cells; (3) CTE-19.20-RG protein and CAR-19 T cells were incubated at 37°C for 10 minutes and then added to target cells. In each case, the target JeKo-1-CD19KO cells were seeded at 1 x 10^4^ cells per well in 50 µl, CTE-19.20-RG was applied in 25 µl medium per well for a final concentration of 1 μg/ml, and CAR-19 T cells were applied to the well as 3:1 or 1:1 ratio to the target cells in 25 µl volume. The plates were incubated at 37°C for 48 hours, then processed for luminescence measurement.

### In vitro experiments modulating antigen density

Expression of CD19 (FMC63-PE, Millipore) and CD20 (CD20-PE, Invitrogen) was evaluated on JeKo-1, Ramos and JeKo-19KO cell lines by flow cytometry. Protein expression was quantified using Bang Bead technology (Bang Labs), and following the manufacturor’s instructions. The cell lines were also evaluated for an increase in apparent CD19 density following incubation with CTE-19.20-RG Engager protein.

in order to assess the additive affect of the CAR-T Engager protein on cytotoxicity, JeKo-1 cells were incubated with CAR-19 T cells at different E:T ratios in the presence or absence of different amounts of CTE-19.20-RG protein. The cells were incubated for 18 hours and then the extent of cytotoxicity was evaluated.

### In vitro model of antigen escape

Wildtype JeKo-1 cells were seeded at 1 × 10^4^ cells in 100 μl per well of complete media in 96 well round bottom plates. CAR-19 T cells in 100 μl complete media were added to give E:T cell ratios of 1:1, 0.3:1, or 0.1:1 and the co-culture was incubated for up to 13 days. Samples were evaluated at days 1, 6, 11 and 13 for the accumulation of a CD19-negative/CD20-positive population, using flow cytometry (FMC63-PE, and anti-CD20-APC, BD). Once a CD19-negative population emerged, the lymphoma cells were counted and replated at 1 × 10^4^ cells in 50 μl complete media. JeKo-1 cells were left untreated in media, or were incubated with 1 × 10^5^ CAR-19 T cells added in 50 μl (E:T ratio of 10:1), with or without CTE19.20-His protein in 50 μl, all in complete media. In a set of control wells the CTE19.20-His protein was added to the JeKo-1 cells alone, without CAR-19 T cells. In each case, the 150 μl co-culture was incubated for 48 hours. Then, the JeKo-1 cells were stained and analyzed by flow cytometry using HIB-CD19-PE (BioLegend) and anti-CD20-APC for cells not incubated with protein or with anti-CD20-APC and anti-ROR1-PE (BioLegend) for samples treated with protein. 7AAD was used to gate out dead cells.

### In vivo models of CD19-negative lymphoma

All *in vivo* protocols were approved by the The Institutional Animal Care & Use Committee (IACUC) of The Cummings School of Veterinary Medicine at Tufts University. To determine the stability of the CD19 knockout phenotype *in vivo*, cells were implanted subcutaneously in NOD-*scid* IL2Rgamma^null^ (NSG) mice and allowed to expand *in situ* for 18 days. The implanted tumors were excised, disaggregated and a single cell suspension was stained with antibodies to human CD19 and CD20.

For tumor efficacy models, 6-10 week old female NOD-SCID/*Il2rg* KO (NSG) mice (Jackson Labs) were acclimated in the vivarium for a minimum of 3 days prior to study initiation. All animals were socially housed in static, sterile bio-contained disposable cages pre-filled with corn cob bedding. Food (LabDiet) and acidified water were provided *ad libitum*. Mice were injected IV with 0.1 mL/animal of 2.5×10^6^ JeKo-19KO cells on day 1. On day 3, all animals were imaged and then randomized to give equivalent average tumor burden in each of 6 cohorts of n = 8 mice/cohort. The following day, different cohorts of mice were given doses of 0, 0.016, 0.08, 0.4 and 2 mg/kg of CTE-19.20, followed by 1 x 10^7^ CAR-19 T cells 3-4 hours later. Protein was then dosed three times weekly for a total of 14 injections. Control cohorts included those treated with CAR-19 cells only and untreated mice.

At least once daily, animals received a cage side health check and clinical observation were performed. Body weights were recorded prior to tumor induction and then three times weekly. Body weights were used to calculate protein and D-luciferin doses. Whole body imaging was performed twice weekly for all animals on study 10-15 minutes after dosing with 15 mg/mL luciferin sc, at 0.2 mL/animal, with isoflurane anesthesia. Lumin values > 1×10^10^ total flux were considered cause for humane euthanasia.

### Pharmacokinetic (PK) measurement by ELISA

Balb/c or NSG mice were dosed once IV with 5 mg/kg CTE-19.20 proteins and blood samples were drawn over time to evaluate protein levels. Serum samples were diluted 10-2000-fold depending on the sample time point. The purified protein standard and diluted serum samples were prepared in the equivilent serum matrix. Samples were analyzed using the FMC63-capture and biotinylated-albumin ELISA described above.

## Results

### Characteristics of Ramos cells, JeKo-1 cells, CD19-deficient JeKo-1 cells and CAR-19 T cells

We evaluated cultured Ramos, JeKo-1 and JeKo-19KO cells for CD19 and CD20 expression. JeKo-19KO cells were further evaluated for CD19 expression after infusion and growth for 18 days in NSG mice. CD19 expression remained deficient in the JeKo-19KO line both *in vitro* and *in vivo*; CD20 expression remained stable (Supplemental Figure 1a-d). Therefore, these cell lines represented CD19-positive and CD19-negative lymphoma, that both express CD20.

CAR-19 T cells were made using T cells from two donors. Exemplary preparations are shown (Supplemental Table 1). In these examples the CAR domain was expressed >80% T cells and robust cytotoxicity was demonstrated at low E:T ratios against the JeKo-1 cell line.

### Characterization of CAR-T Engager (CTE) proteins

CTE proteins bridge the CAR-T cells to new antigens via antigen binding domains derived from antibodies and related scaffolds (18–20). We created a CTE protein with an anti-CD20 llama VHH linked to a stabilized form of CD19 ECD, followed by an anti-albumin VHH. The apparent binding affinity, EC50, of the anti-CD20 VHH for human CD20 was 4.2 and 2.8 pM in the quantitative flow cytometry assay using JeKo-1 cells and JeKo-19KO cells, respectively (Supplemental Figure 1e). CTE-19.20 proteins were produced in two transient transfections. A C-terminal His-tagged form called CTE-19.20-His was produced from Expi293 cells (0.1g/L yield, 100% purity) and CTE-19.20-RG was made from CHO cells (1g/L, 98.7% purity). CTE-19.20-RG is identical to CTE-19.20-His, minus the 6x His-tag. The purified proteins showed similar bands by PAGE analyses and showed high purity with minimal aggregation in SEC traces (Supplemental Figure 2a-c). The expected molecular weights of CTE-19.20-His and CTE-19.20-RG were 56.4 and 55.6 kDa respectively, however the proteins ran at higher molecular weights on PAGE gels, presumably due to N-linked protein glycosylation.

CTE proteins were evaluated for binding to the three target molecules: CD20, albumin, and the anti-CD19 MAb FMC63. Anti-CD19 was used to capture the CTE-19.20 proteins in ELISA assays, followed by incubations with biotinylated human albumin or human CD20, and the detection reagents. Concentration curves were generated for each protein in each ELISA format (Figure 1a,b), and the binding affinities were calculated from multiple repetitions of the assays. The two protein preparations had very similar EC50 values of ∼ 20 ng/ml (Table 1).

**Figure 1.**
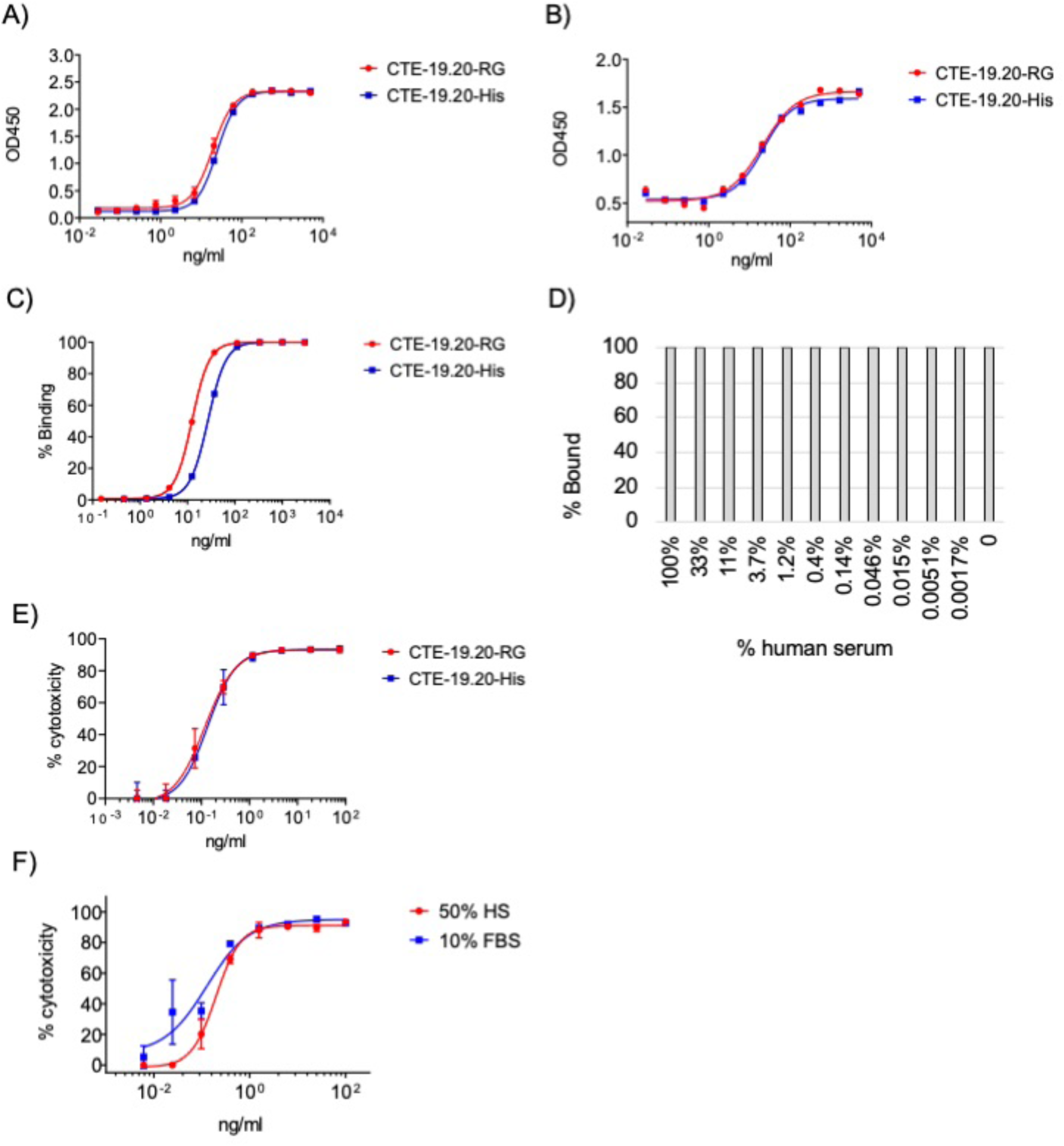
CTE proteins bind to anti-CD19 antibody coated plates and can be detected by ELISA assay using A) biotinylated albumin, and B) biotinylated CD20 membranes. C) Binding of CTE-19.20 proteins to JeKo-19KO cells. D) Binding of CTE-19.20 proteins to JeKo-19KO cells in the presence of excess serum. E) Dose dependent cytotoxicity mediated by CTE-19.20 proteins in the presence of JeKo-1KO cells and CAR-19 T cells (54% CAR-positive). F) CTE-19.20-RG-mediated cytotoxicity in the presence of media (10% FBS) or media further supplemented with human serum (50% HS), and CAR-19 T cells (78% CAR-positive).

**Table 1.** Characterization of CTE-19.20 protein activities. Top: apparent binding affinities in two ELISA assays. Middle: apparent binding affinity of binding to JeKo-19KO cells as measured by flow cytometry using anti-CD19 antibody FMC63 detection. Bottom: IC_50_ values derived from the cytotoxicity assay using CAR-19 T cells (81% CAR-positive), JeKo-19KO cells and CTE-19.20 proteins. The activities of the two CTE-19.20 preparations were evaluated by unpaired, two-tailed, T-test: *, p=0.011.

| Protein | Anti-CD19 capture /<br>CD20 detection: EC<sub>50</sub> (ng/ml), sd (n) | Anti-CD19 capture /<br>Albumin detection: EC<sub>50</sub> (ng/ml), sd (n) |
| --- | --- | --- |
| CTE-19.20-His | 25.4, 5.0 (n=2) | 26.7, 6.5 (n=6) |
| CTE-19.20-RG | 16.4, 2.4 (n=3) | 23.6, 7.4 (n=7) |
| Protein | Flow cytometry binding affinity EC<sub>50</sub> (ng/ml), sd (n) |  |
| CTE-19.20-His | 26.6, 2.2 (n=3)* |  |
| CTE-19.20-RG | 11.9, 2.3 (n=3) |  |
| Protein | Cytotoxicity IC<sub>50</sub> - mean (ng/ml), sd (n) |  |
| CTE-19.20-His | 0.14, 0.12 (n=3) |  |
| CTE-19.20-RG | 0.13, 0.09 (n=11) |  |

Using JeKo-19KO cells that lack CD19, we evaluated the affinity of protein binding to cell surface expressed CD20, using dose titrations. For this flow cytometry assay, cells were incubated with proteins and then with fluorescently labeled anti-CD19 MAb. Dose response binding curves were generated (Figure 1c). The CTE-19.20 proteins bound to JeKo-19KO cells with similar EC50 values, ∼20 ng/ml, calculated from multiple repetitions of the assay (Table 1).

These results showed that CTE-19.20 proteins bound their targets with high affinity in both plate-based ELISA and cell-based flow cytometry formats. We further demonstrated that binding to albumin in up to 100% human serum did not interfere with CTE-19.20-RG binding to CD20 or to anti-CD19 as determined by percent binding in the flow cytometry assay (Figure 1d), however in the presence of 33%-100% human serum the mean fluorescence intensity (MFI) was reduced by ∼30% To extend the binding results to a functional assay we performed cytotoxicity assays using JeKo-19KO cells. Dose response binding curves were generated from cytotoxicity assays. Representative results are shown (Figure 1e), and the mean IC50 values (∼0.13 ng/ml) from multiple experiments are shown in Table 1. Serum albumin did not impact the degree of cytotoxicity (Figure 1f). This shows that the modest reduction in MFI did not affect cytotoxicity.

CTE-19.20 proteins therefore potently redirected CAR-19 T cells to kill CD19-negative lymphoma cells. Next it was important to show that the presence of the CTE protein did not effect direct CAR-19 cytotoxicity, ie. to lymphoma cells expressing CD19. Wildtype JeKo-1 cells were cultured with CAR-19 T cells alone or with up to 1 μg/ml CTE-19.20-RG, a saturating amount of protein, plus CAR-19 T cells. There was no impact of CAR-T Engager protein on CAR-19 cytotoxicity as compared to culture of the target lymphoma cells with CAR-19 alone, at the 3:1 and 1:1 E:T ratios (Supplemental Figure 3a,b).

We next evaluated whether CTE-19.20-His protein could induce internalization or downregulation of cell surface CD20, as measured by protein binding. JeKo-19KO cells were incubated at 4°C or 37°C to examine change in the total percent of cells binding to CTE-19.20 and the mean fluorescent intensity (MFI) of binding. There was no change in percent cells bound at 4°C, a temperature at which most proteins will not alter surface expression, nor was there any change at 37°C (Supplemental Figure 4a). This indicated that no CD20 downregulation or internalization was occurring. Indeed, the MFI of CTE-19.20-His staining increased and was maintained through 6 hours at 37°C (Supplemental Figure 4b).

To extend the CTE protein saturation and internalization results, we tested whether the order of addition of CTE protein to the cytotoxicity assay impacted cytotoxicity. We evaluated whether preincubation of 1 μg/ml CTE-19.20-RG with CAR-19 T cells, or with target tumor cells, would impact cytotoxicity, compared to adding the components simultaneously. Two E:T ratios of CAR-19 T cells to JeKo-19KO cells were used, 3:1 and 1:1, and extent of cytotoxicity was compared to JeKo-19KO cells plus the CAR-19 T cells or to the JeKo-19KO cells plus protein (Supplemental Figure 5a,b). There was no difference in the extent of cytotoxicity regardless of the order in which the components were preincubated. This suggests a) that pre-binding of the CTE-19.20 protein to CAR-19 T cells does not impact CAR function and b) that the cytotoxicity triggered by binding of CAR-19 to target JeKo-19KO cells is not competed by a saturating concentration of CTE-19.20-RG protein.

### Evaluation of CD19 antigen density and enhanced cytotoxicity in the presence of CTE-19.20 protein

We used a fluorescent bead assay to quantify the increase of CD19 antigen density on the cell surface as increasing amounts of CTE-19.20-RG protein were incubated with the JeKo-19KO cells (Figure 2a). The background (no protein added) was equivilent to 515 CD19 copies. With increasing concentrations of CTE-19.20-RG, the apparent CD19 density increased until it was very similar to the number of CD20 molecules present: approximately 263,000 CD19 copies and approximately 267,000 CD20 copies (Figure 2a).

**Figure 2.**
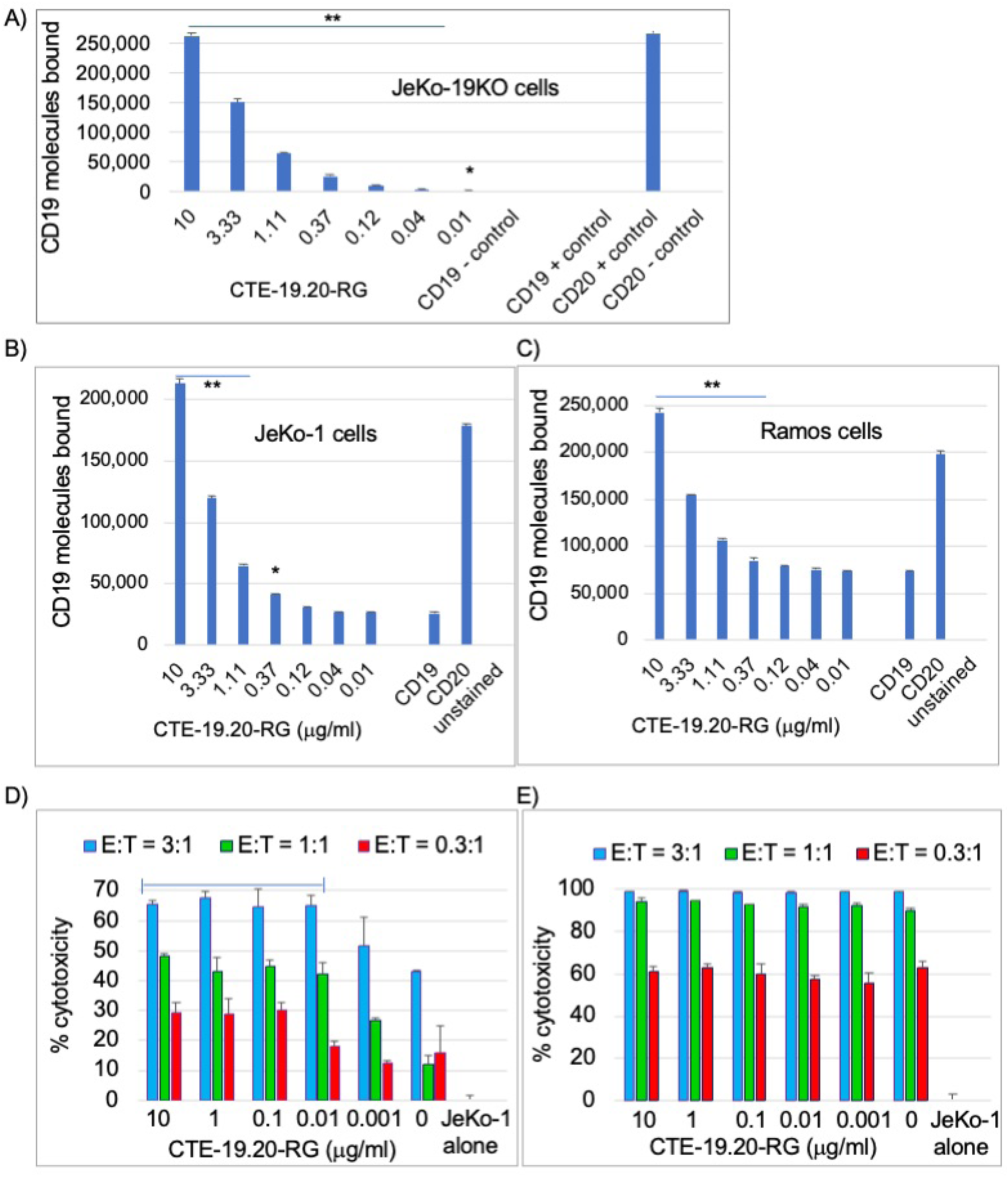
Characterization of antigen density and cytotoxicity. A-C) Increase in apparent CD19 density on (A) JeKo-19KO, (B) wildtype JeKo-1 and (C) Ramos cells in the presence of increasing concentrations of CTE-19.20-RG protein vs CD19 staining and CD20 staining alone. D,E) Cytotoxicity of CAR-19 T cells (77% CAR-positive) against JeKo-1 cells at different E:T ratios in the presence on increasing concentrations of CTE-19.20-RG after (D) 18 hours and (E) 48 hours. Data were analyzed by one-tailed T test, *: p < 0.05, **: p < 0.02 vs CD19 controls (A-C) and p < 0.02 vs ‘0’ (D).

JeKo-1 and Ramos cells express both CD19 and CD20. As is typical of lymphoma, CD20 was brighter than CD19 on both cell lines as measured by flow cytometry, with a fluorescent intensity approximately 5 times high on JeKo-1 cells and 2-3 times higher on Ramos cells (Supplemental Figure 1a). On the JeKo-1 cell line, approximately 25,500 copies of the CD19 protein were detected, and approximately 177,000 copies of the CD20 protein (Figure 2b).

Similarly, on Ramos cells, approximately 74,000 copies of the CD19 protein were detected, and approximately 198,000 copies of the CD20 protein (Figure 2c). CD19 density was measured as increasing amounts of CTE-19.20-RG protein was incubated with the cells. In both cell lines, incubation with > 120 ng/ml of the Engager protein led to an increase in the apparent CD19 density, up to 10 μg/ml, at which point the signal detected was similar to that combination of the CD19 and CD20 antigen densities on each cell type, thus from ∼25,000 copies to ∼220,000 copies of CD19 on JeKo-1 cells, and from ∼74,000 copies to ∼240,000 copies of CD19 on Ramos cells (Figure 2b,c).

Increased antigen density should enhance CAR-19 mediated cytotoxicity against cells that naturally express CD19. We tested this hypothesis by incubating JeKo-1 cells with increasing amounts of CTE-19.20-RG protein and evaluating cell killing at 18 hours. There was a nominal, statistically insignificant increase when the E:T ratio was set to 3:1, but a marked increase at the lower E:T ratios of 1:1 and 0.3:1 (Figure 2c). Thus, in the initial period of cytotoxicity and at low E:T ratios, the increased apparent CD19 antigen density impacted the efficiency of CAR-19 cytotoxicity (Figure 2d). The effect was absent after 48 hours at which point the extent of killing had been maximized (Figure 2e).

### In vitro models of lymphoma antigen loss and escape from therapy

Previously we showed that CAR-19 T cells can target and kill CD19-negative cells via a CD19 ECD linked to the anti-CD20 antibody leu16 scFv (20). Here we used CTE proteins and several cell models to extend those findings. CTE-19.20 proteins bound to JeKo-19KO cells, a CD19-negative lymphoma, and dose responsive cytotoxicity was induced when CAR-19 T cells were added (Figure 1e). Table 1 shows that the two CTE-19.20 protein preparations were equally potent in inducing cytotoxicity.

JeKo-1 cells have a very high proliferative index and we hypothesized that they could provide a useful model of antigen escape from CAR-T cell therapy. Therefore, we characterized the phenotype of JeKo-1 cells under CAR-T pressure. JeKo-1 cells are CD19-positive and CD20-positive and the vast majority are double-positive (Figure 3a). We derived an antigen escape model using various E:T ratios whereby the number of CAR-T cells was varied versus a fixed number of JeKo-1 cells and by varying the length of time in culture (Figure 3b-d).

**Figure 3.**
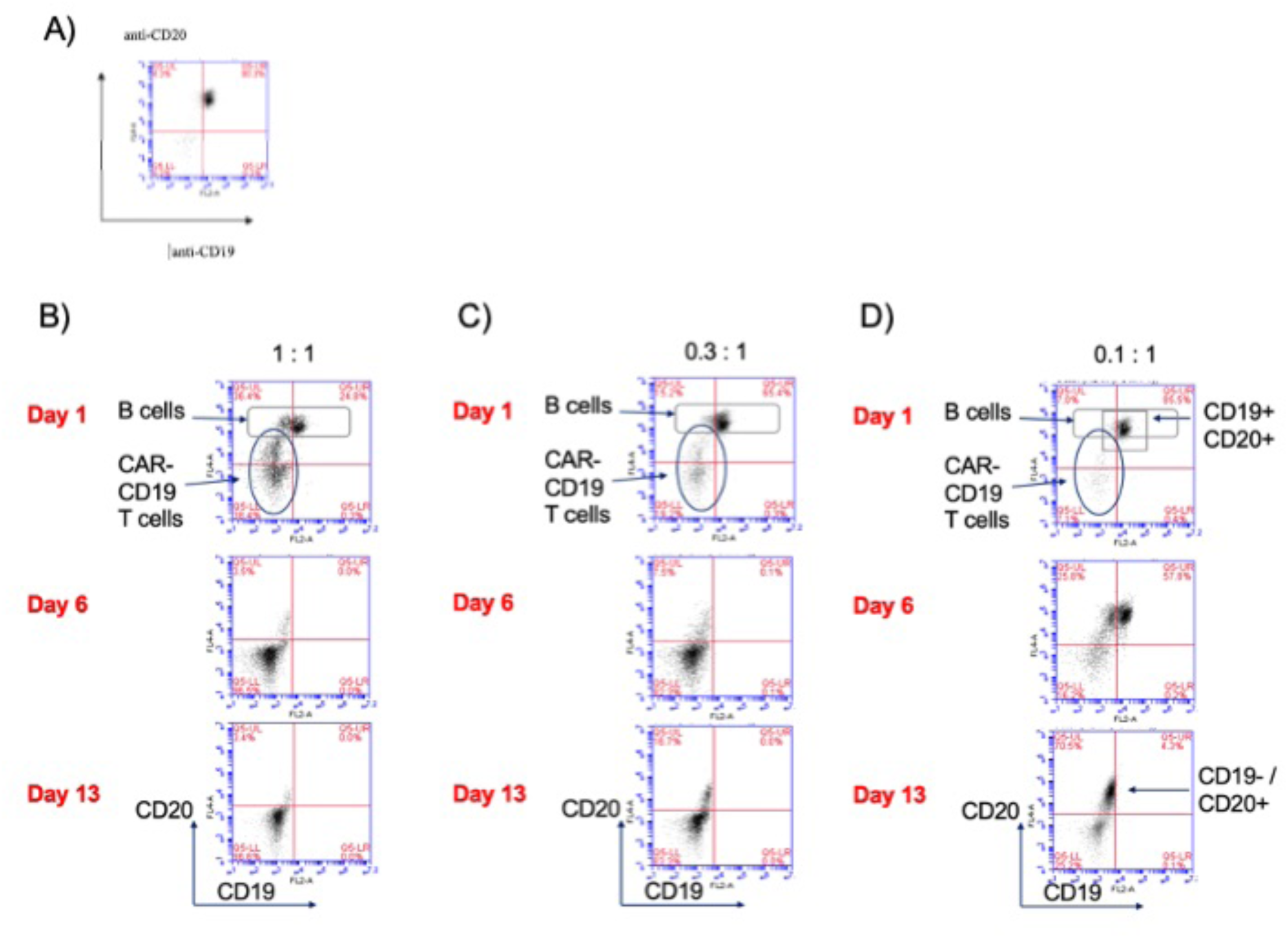
Development of a JeKo-1 antigen escape model from treatment with CAR-19 T cells (55% CAR-positive) *in vitro.* The position of CAR-19 cells and the target JeKo-1 cells is indicated in the flow cytometry profiles is shown to the left of the panels. The designation of CD19-positive and CD20-positive cell populations is shown to the right of the panels. A) JeKo-1 cells express CD19 and CD20, B) E:T ratio 1:1 from day 1 – 13. C) E:T ratio 0.3:1 from day 1 – 13. D) E:T ratio 0.1:1 from day 1 – 13.

JeKo-1 cells escaped from CAR-19 cytotoxicity via cell population-level loss of CD19 expression (Figure 3). This occured even at a 1:1 E:T ratio as seen in column B, days 6 and 13. The effect was more pronounced at the 0.3:1 and 0.1:1 E:T ratios as seen in column C, days 6 and 13, and column D, where the effect was especially pronounced at day 13. Notably, the 0.1:1 E:T ratio was ineffective in inducing cytotoxicity (see Figure 2, Day 6) yet even this modest level of selective pressure imposed by the CAR induced a shift from a mainly double-positive (CD19 and CD20 positive) population on day 1 to a mixed population on day 6 to a population dominated by CD19-negative, CD20-positive cells by day 13.

Since antigen loss and tumor cell escape occurred across this range of E:T ratios and days of incubation, a second study evaluated the effect of adding the CTE-19.20 protein to the culture. As a control, the lymphoma cells were replated without the addition of either CAR-19 T cells or CTE protein. After 48 hours, a mixed phenotype lymphoma cell population had emerged consisting of CD20-positive/CD19-positive cells, CD20-positive/CD19-low cells, and CD20-positive/CD19-negative cells (Figure 4a). The addition of CAR-19 T cells plus CTE-19.20 protein, but not the CAR-19 T cells alone nor the CTE-19.20 protein alone, was sufficient to kill all target JeKo-1 cells post escape and recovery (Figure 4b). These results show that lymphoma cell escape by loss of CD19 expression can be rapidly induced *in vitro* and fully reversed using the CD20-targeting bridging protein, CTE-19.20 and CAR-19 T cells.

**Figure 4.**
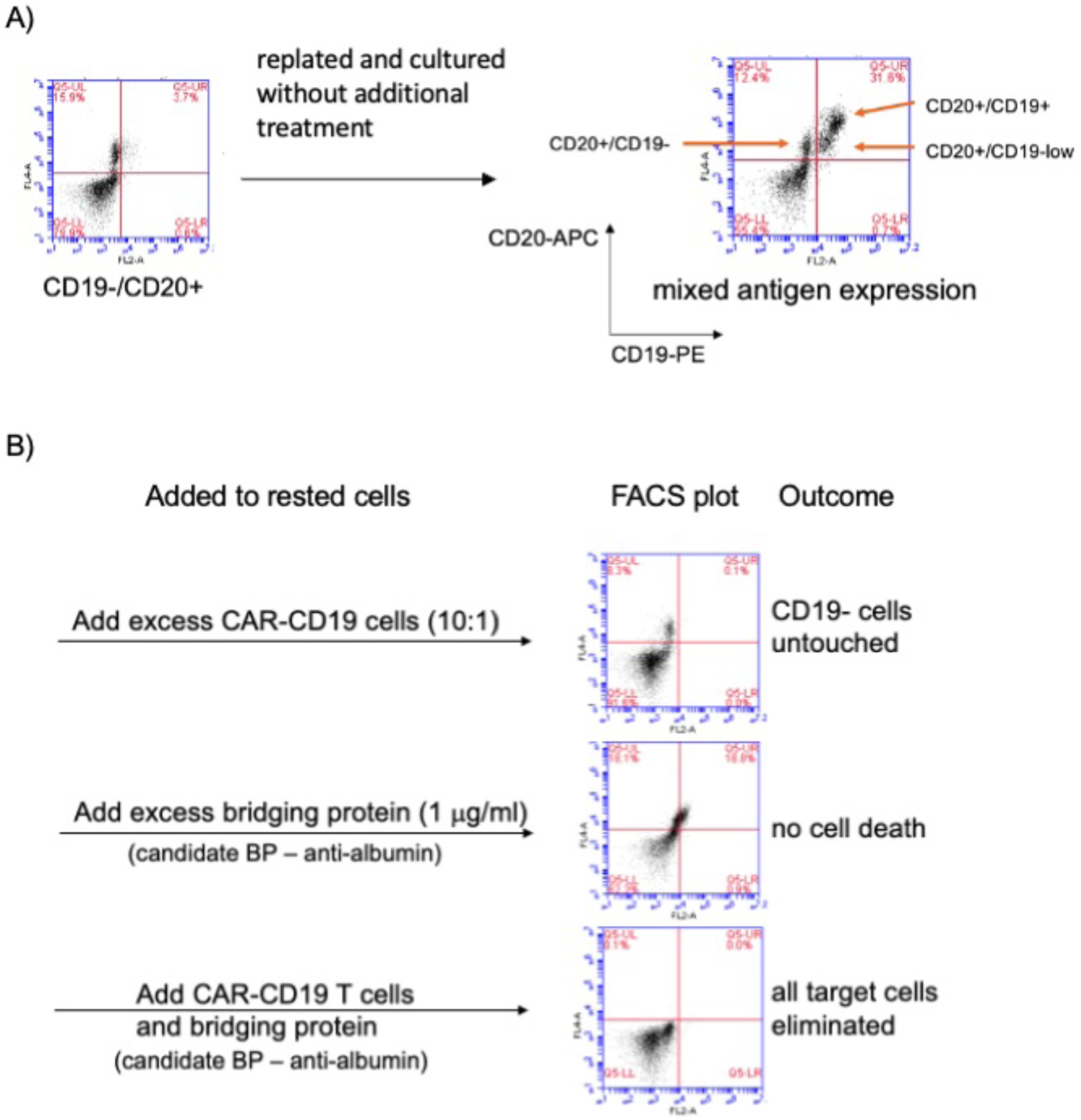
Reversal of antigen-loss mediated relapse from CAR-19 treatment *in vitro*. A) CAR-19 cells (55% CAR-positive) were incubated with JeKo-1 cells (E:T 1:1) for 13 days, and then rested for 48 hours. Expression of CD19 and CD20 was monitored by FACS. B) CTE-19.20 protein was added to the rested culture, or CAR-19 T cells were added to the rested culture, or both CTE-19.20 and CAR-19 T cells were added to the rested culture.

### In vivo modeling

We used the JeKo-19KO cell line, CTE19.20 proteins and CAR-19 T cells in several *in vivo* models. NSG mice were injected with JeKo-19KO lymphoma cells which were implanted for 4 days prior to treatment with CAR-19 T cells and CTE-19.20 protein. CAR-19 T cells were injected once, and CTE-19.20 protein was given 3x weekly. Tumor burden was quantified using luciferase-generated luminescence.

In the initial study the CTE.19.20-His and CTE-19.20-RG protein preparations were screened for *in vivo* activity at a dose of 2 mg/kg in the presence of CAR-19 T cells as compared to cohorts given CAR-19 T cells only, or left untreated. Thrice weekly dosing with CTE-19.20 proteins restrained lymphoma development through day 31, at which time the untreated mice and the CAR-19 treated mice began succumbing to disease (Figure 5a).

**Figure 5.**
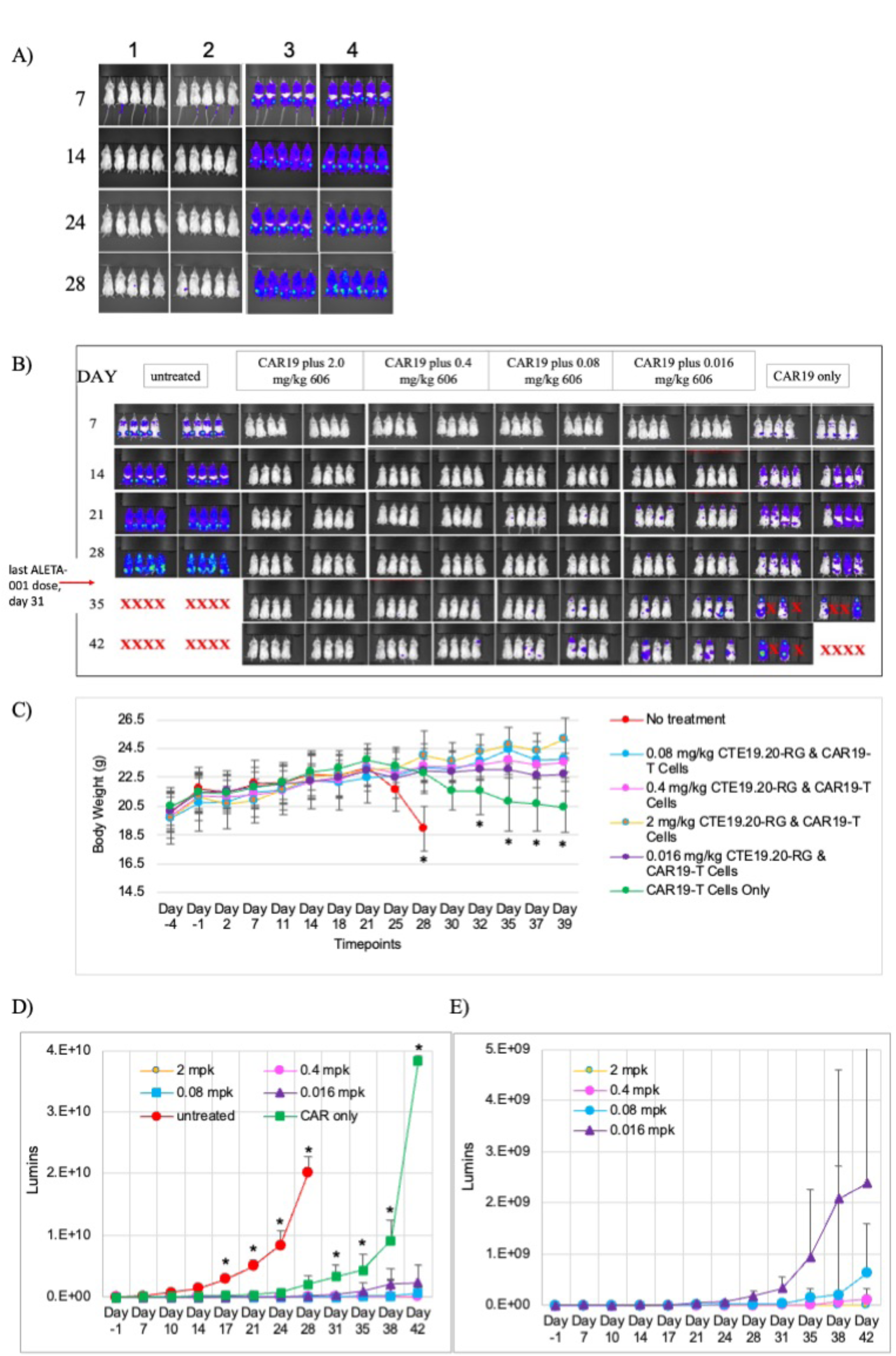
CTE-19.20 proteins given with CAR-19 T cells successfully treat lethal CD19-negative lymphoma *in vivo*. A) Comparison of the efficacy of the 2 mg/kg dose of CTE-19.20-His (1) and CTE-19.20-RG (2) with CAR-19 cells alone (3) and donor-match UTD cells (4). B) Dose response and efficacy of treatment with CTE-19.20-RG, during and after the dosing period. C) CTE-19.20-RG treatment supports continued weight gain in animals challenged with otherwise lethal lymphoma. D) CTE-19.20-RG treatment significantly reduces tumorburden as assessed by JeKo-19KO luciferase activity. E) Luciferase activity of the treatment groups only, graphed to demonstrate dose-responsive efficacy. *: p < 0.02 by two-way T test of control cohorts vs all treatment cohorts.

Next we performed a dose titration of CTE19.20-RG across 4 cohorts: 2 mg/kg, 0.4 mg/kg, 0.08 mg/kg and 0.016 mg/kg. Animals were dosed with protein until day 31 at which time dosing was stopped in order to evaluate lymphoma relapse. Nine days later, on day 42, there was no sign of luminescence in animals treated with CAR-19 plus 2 mg/kg CTE-19.20-RG and only a minimal signal was seen in 2 of 8 animals treated with CAR-19 plus 0.4 mg/kg CTE-19.20-RG). Lower doses of CTE-19.20-RG were less effective in preventing relapse after dosing cessation (Figure 5b).

Body weight is a quantitative means of tracking animal health and is used in conjunction with clinical examinations to ensure humane euthanasia in cancer models. Mice treated with CAR-19 plus CTE-19.20-RG protein continued to gain body weight throughout the protein dosing period (until day 31), while the control animals began losing weight at approximately day 25 (Figure 5c). Even after dosing with CTE-19.20 protein stopped, the animals in the treated cohorts maintained body weight through the end of the study (day 42) and the highest dose-treated cohort, that had received 2 mg/kg of CTE-19.20 up until day 31, continued to gain weight with the last recorded weight taken on day 39 (Figure 5c).

Lumin intensity was graphed for each cohort, showing a rapid increase in the untreated group and a slightly delayed but still rapid increase in the CAR-19 treated group. The termination of the lines for the untreated and CAR-19-treated groups shows when all animals in those cohorts succumbed to disease (Figure 5d). To discern differences between CTE-19.20-RG dose cohorts we removed the control groups from the graph (Figure 5e). A clear dose response then became evident, showing a complete lack of luminescent signal in the 2 mg/kg CTE-19.20 cohort through day 42 (Figure 5e). Since the last protein dose was given on day 31, this suggests that these animals had eliminated the CD19-negative lymphoma.

Protein efficacy *in vivo* is a function of the pharmacokinetic (PK) properties of the injected biologic. We evaluated the PK properties of CTE-19.20-His and CTE-19.20-RG in mice. The half-life of the biologic in mouse depends on albumin-mediated recirculation via FcRN binding. The anti-CD20 domain used in the CTE-19.20 proteins does not bind mouse CD20 and but the anti-albumin domain binds to mouse albumin weakly with an apparent EC50 of 177 nM (Supplemental Figure 6). The human CD19 ECD has no known ligands other than the B cell transmembrane proteins CD21 and CD81, with which it forms a compex *in cis* on the surface of B cells (22).

The CTE-19.20-His protein was evaluated in both Balb/c and NSG mice after a single injection IV and showed a similar half-life of 21 hours in both strains (Table 2). However, we sampled NSG mice 24 hours after injection of CTE-19.20-RG and could not detect the protein in mouse serum. Therefore, we performed a more time-point intensive analysis of CTE-19.20-RG PK in Balb/c mice. Surprisingly, the half-life of CTE-19.20-RG in mouse was shorter than expected, at 7.6 hours (Table 2).

**Table 2.** Evaluation of CTE-19.20 protein half-life in mice and the degree of protein sialylation.

| **Protein** | **T<sub>1/2</sub>, Balb/c** | **T<sub>1/2</sub>, NSG** | **cIEF, % basic**<br>**pH peaks** | **cIEF, % acidic**<br>**pH peaks** |
| --- | --- | --- | --- | --- |
| CTE-19.20-His | 20.63 | 27.7 | 26.2 | 73.8 |
| CTE-19.20-RG | 7.6 | nd | 100 | 0 |

Protein half-life can also be impacted by sialylation, the post-translational, covalent addition of terminal sialic acid to glycosylated proteins (23). Proteins with more sialyation have longer half-lives *in vivo* (24, 25). We used capillary isoelectric focusing (cIEF) separation to evaluate CTE-19.20 protein sialylation. In cIEF, a continuous pH gradient is formed by applying voltage across a capillary filled with carrier ampholytes, and this separates proteins by their isoelectric point (pI). Simplistically, proteins can be evaluated as the percent of basic and acidic peaks, where acidic peaks represent sialylated protein. We found that while the CTE-19.20-His protein contained a mixture of acidic and basic peaks, the CTE-19.20-RG protein was almost wholly basic, indicating minimal sialylation (Table 2). These results may explain the differential PK properties of the two protein preparations.

## Discussion

We previously described a novel approach to the extend the reach of CAR therapies using diverse CAR-T Engager (CTE) proteins that contain the CD19 ECD linked to an antigen binding domain, such as an scFv or llama VHH (18–20). These CD19-based CTE proteins bind to any antigen targeted, coat the desired tumor with CD19, and retarget CAR-19 T cells to kill those tumor cells via the bound protein (18–20). One therapeutic focus has been a modality in which the CAR T cell secretes the CTE protein (18, 19). This is thematically similar to the secretion of bispecific CD3-based T cell engagers from CAR T cells (26, 27), with the additional characteristic of CAR domain specificity. We validated the approach preclinically using both solid and hematologic tumors, showing *in vitro* and *in vivo* efficacy against Her2-positive ovarian carcinoma and CLEC12A-positive acute myeloid leukemia cells (18, 19).

The use of CTE proteins as injectable biologics provides an alternative therapeutic modality. We have developed a CTE biologic to prevent and reverse relapses from CAR-19 cell therapy for B cell lymphoma and leukemia. This novel CTE contains a CD19 ECD that binds to anti-CD19 domains expressed by CAR-19 T cells. Two additional domains were linked to the CD19 ECD, an anti-CD20 VHH and an anti-albumin VHH, both derived from immunized llamas. The resulting protein, CTE-19.20, functions by binding to both CD20 on target tumor cells and to CAR-19 T cells, thereby triggering cytotoxicity. A favorable circulating half-life is engineered via binding to albumin which binds the FcRN receptor to slow protein degradation (28). Our preliminary work suggests that CTE protein sialylation may also contribute to the *in vivo* circulating half-life in mouse models.

CAR-19 T cells can bind directly to CD19 in the presence of the CTE-19.20 protein, and gain the ability to bind the CD19 ECD that is bound, via the anti-CD20 VHH, to CD20. This additional source of CD19, coated on CD20 on the cell surface, increases available CD19 antigen density. We quantified the increase in CD19 density on JeKo-19KO, JeKo-1 and RAMOS cells incubated with CTE-19.20-RG protein, showing that the signal increased to match the available number of CD20 molecules to which the CAR T engager could bind (Figure 2). Importantly, lymphoma cells often have high cell surface expression of CD20 (29), providing for abundant CTE binding. One caveat of the “bang bead” analysis was the relative insensitivity of the assay when adding CTE-19.20 protein to CD19-expressing cells, such that the increase in antigen density was detectable only at concentrations of CTE-19.20-RG protein that are higher than the cytotoxicity IC50. This phenomena is also noted in our flow cytometry analyses, where the binding EC50 measured by flow cytometry is always much higher than the cytotoxic IC50 (see Table 1). This difference indicates only a subset of CAR domains need to be bound in order to trigger cytotoxicity. By adding the CTE-19.20 protein to increase the available antigen present on the target lymphoma cell line, we triggered the critical threshold for cytotoxic activation of the CAR-T cell more quickly, and at lower E:T ratios.

There is widespread appreciation that antigen density and antigen loss in response to therapy are key drivers of successful CAR-19 T cell treatment (30). Low antigen density or antigen loss uncouples the CAR-T cells from targets, leading to loss of response. Since CTE19.20 protein creates a CD19 ‘depot’ that is bound to CD20 and independent of cellular CD19 expression, this solution should diminish the rate of relapse due to antigen loss, as we modeled using the *in vitro* “antigen escape” assays (Figures 3 and 4). Indeed, the loss of CAR activity quickly led to the re-emergence of CD19-positive cancer cells (Figure 4), illustrating the power of natural selection on a population of rapidly dividing cancer cells. At low E:T ratios, and over several weeks of culture, a CD19-negative cell population emerged (Figure 3). Once removed from the selection pressure imposed by CAR-19 T cells, a mixture of CD19-negative, CD19-dim and CD19-bright cells was observed that could then be eliminated by the addition of CAR-19 T cells and CTE-19.20 protein (Figure 4). The transient nature of CD19 loss provides preclinical evidence for the hypothesis that at least some of the “CD19-positive” relapses recorded in the clinical literature may actually result from transient loss and then rapid recovery of CD19 expression, with the CAR-19 T cell pool collapsing in between; this would be difficult to discern without longitudinal analyses (14). Similar phenomena have been documented in the setting of multiple myeloma treatment with anti-BCMA CAR therapy (12).

Detailed analyses of CD19 antigen density required for productive CAR-19 persistence and function identified specific expression thresholds below which CAR-T cells lost functionality (13, 27). Antigen density as measured by immuno-histochemistry was not sensitive enough to distinguish tumor populations with cells above or below those thesholds, whereas quantitative flow cytometry was sufficiently sensitive (13, 27). The importance of antigen density as a driver of CAR-T cell activity has also been modeled with anti-CD22 CAR-T cells (32) and demonstrated by the use of γ-secretase-antagonism to prevent BCMA shedding by multiple myeloma cells in response to anti-BCMA CAR-T therapy (33). Antigen density thresholds have been identified in preclinical studies of CAR-T cells directed against diverse antigens including CD22, CD38, ALK and CD20 (32, 34–36).

Clinically, antigen density associated with therapeutic response has been described for CAR-19 and CAR-BCMA therapies for B cell lymphoma and multiple myeloma (15, 31, 33, 37, 38). For example, CD19 antigen loss was associated with relapse from lymphoma in 62% of patients treated with an anti-CD19 CAR-T therapy, as assessed using quantitative flow cytometric methods (15, 31). Many mechanisms for CD19 antigen loss have been described, including transcriptional downregulation of expression and genetic alterations that prevent expression (13, 30, 39, 40).

Clinical trials in NHL, ALL and CLL have consistently demonstrated key features underlying patient outcome, leading to rapid relapse or, conversely, durable remission. Prominent among the features that predict successful patient outcome is rapid and prolonged expansion of CAR-T cells in the first 0-90 days post infusion, measured, for example, as CAR-T AUC0-28 or AUC0-48 (10,28,29). Conversely, patients who have lower CAR-T cell expansion, for a shorter time, typically do not achieve a durable response (43–45). Increased CD19 antigen density enabled more rapid and efficient cytotoxicity with low levels of CAR-T cells relative to targeted wildtype JeKo-1 cells (Figure 2d). The increase in antigen density when incubating lymphoma cells with a CTE-19.20 protein was most apparent when the density of CD20 on the target cell surface was much higher than the CD19 density. This was clearly demonstrated using the JeKo-1 MCL cell line (Figure 2b). Even with a smaller difference in expression of CD20 and CD19, as in the Ramos Hodgkin lymphoma line, additive CD19 density was observed (Figure 2c). Of note, CD20 expression is higher than CD19 expression in most lymphoma although expression of both markers is variable (29, 46). CD20 expression is also variable in CLL and is absent in some ALL patients (47). Across these indications, in patients with CD20-positive malignancies, the ability to increase the apparent antigen density of CD19 would be expected to improve the response of CAR-19 T cells (31).

Our *in vivo* modeling used CD19-deficient cells to represent relapse from CAR-19 cell therapy. *In vivo*, CAR-19 cells were unable to detect and kill the JeKo-19KO cells (Figure 5a), leading to systemic and lethal lymphoma. Administration of CTE-19.20-RG protein along with the CAR-19 T cells prevented lethality. Of note, upon cessation of dosing with CTE-19.20-RG protein, animals remained lymphoma free for 11 days, at which time all control animals had expired (Figure 5b). On day 42, when the study ended, 75% of animals in the three highest dose cohorts had no luminescent signal (Figure 5e) and only 1 of 32 treated animals, from the lowest dose cohort, had expired. In contrast, 14/16 animals in the control cohorts had expired. This result was achieved despite the post-study finding that the CTE-19.20-RG protein had an unexpectedly short half-life in mouse of approximately 7.5 hours. The half-life is correlated with the extent of sialylation (Table 2), and current manufacturing and process development is focused on producing highly sialylated protein.

Patient relapse from CAR-19 therapy occurs rapidly, often within the first months following therapy (9–11). The kinetics of relapse offer the potential to intervene using CTE-19.20 protein, using several different clinical designs. In the first instance we are developing this molecule with the goal of treating patients who have already received a CAR-19 therapy, by evaluating their clinical response through the first few months post CAR infusion. Diverse biomarkers can be used to track response and risk of relapse. PET imaging and ctDNA analyses have emerged as sensitive means of tracking lymphoma and leukemia regression (48–50). Q-PCR or deep sequencing methods have provided a picture of CAR-19 expansion and persistence over the first critical weeks and months after treatment (51). Flow cytometry can track T cell expansion and also antigen expression on the targeted cancer cells (51–54). Using such methods, patients at risk of relapse could be identified and treated with the CTE-19.20 biologic to improve outcomes. Pre-clinical proof of principal for CAR reactivation via vaccination suggests this is a productive approach (55). Clinical examples include the response of CAR-19 T cell populations to checkpoint inhibition (56, 57). The CTE biologic can also be delivered in combination with CAR-19 T cells initially, to simultaneously target the two antigens, CD19 and CD20.

There are multiple efforts to address antigen loss, including dual-CARs and TanCARs targeting combinations of CD19, CD20, or CD22, for example (58). Nevertheless, combination of CAR-CD19s with the CTE biologic has the immediate advantage of synergy with marketed CAR-19 therapies, with the potential to reduce relapses significantly. We note that other factors impact patient responsiveness to CAR-19 treatment. Suboptimal CAR-T cell expansion can occur if the patient T cell pool has intrinsically limited proliferative potential, as has been described after multiple rounds of aggressive chemotherapy as standard-of-care prior to cell therapy (59). Limited response to CAR-T cell therapy has also been associated with immunosuppression, for example in the lymphoma tumor microenvironment (56).

In summary, we have described an optimal and straightforward approach to productive and sustained activation of CAR-19 T cells, using a potent CD19-anti-CD20 bridging protein with an extended half-life. We note that the extension of our approach to other CAR-T cell relapse settings is feasible, and we have built BCMA-containing CTE proteins capable of killing BCMA-negative cell lines. The use of the CTE biologic approach is in fact viable for any single antigen-targeting CAR-T cell, where relapse and resistance through antigen loss or modulation is almost inevitable in a large subset of treated patients.

## Acknowledgements

Alpha Preclinical LLC, Grafton MA USA performed the *in vivo* studies. Alyssa Birt and Fay Dufort produced some of the expression constructs, lentiviral vector particles and CAR T cells used in this study. Donna Grant provided expert guidance for the PK analysis. Victoria Arena De Souza provided a critical review of the manuscript prior to submission.

## Data availability

The authors confirm that the data supporting the findings of this study are available within the article and its supplementary materials, and that raw data generated at Contract Research Organizations that support the findings of this study are available from the corresponding author [PDR] on request.

## References

1. Ruella M, June CH. Chimeric Antigen Receptor T cells for B Cell Neoplasms: Choose the Right CAR for You. Curr Hematol Malig Rep 2016;11(5):368–384.

2. Chong EA, Ruella M, Schuster SJ. Five-Year Outcomes for Refractory B-Cell Lymphomas with CAR T-Cell Therapy. New England Journal of Medicine 2021;384(7):673–674.

3. Neelapu SS et al. Long-Term Follow-up Analysis of ZUMA-5: A Phase 2 Study of Axicabtagene Ciloleucel (Axi-Cel) in Patients with Relapsed/Refractory (R/R) Indolent Non-Hodgkin Lymphoma (iNHL). Blood 2021;138(Supplement 1):93.

4. Cappell KM et al. Long-Term Follow-Up of Anti-CD19 Chimeric Antigen Receptor T-Cell Therapy. J Clin Oncol 2020;38(32):3805–3815.

5. Schuster SJ et al. Long-Term Follow-up of Tisagenlecleucel in Adult Patients with Relapsed or Refractory Diffuse Large B-Cell Lymphoma: Updated Analysis of Juliet Study. Biology of Blood and Marrow Transplantation 2019;25(3, Supplement):S20–S21.

6. Grupp SA et al. Tisagenlecleucel for the Treatment of Pediatric and Young Adult Patients with Relapsed/Refractory Acute Lymphoblastic Leukemia: Updated Analysis of the ELIANA Clinical Trial. Biology of Blood and Marrow Transplantation 2019;25(3):S126–S127.

7. Lin JK et al. Cost Effectiveness of Chimeric Antigen Receptor T-Cell Therapy in Multiply Relapsed or Refractory Adult Large B-Cell Lymphoma. J Clin Oncol 2019;37(24):2105–2119.

8. Brudno JN, Kochenderfer JN. Recent advances in CAR T-cell toxicity: Mechanisms, manifestations and management. Blood Rev 2019;34:45–55.

9. Maude SL et al. Tisagenlecleucel in Children and Young Adults with B-Cell Lymphoblastic Leukemia. New England Journal of Medicine 2018;378(5):439–448.

10. Neelapu SS et al. Axicabtagene Ciloleucel CAR T-Cell Therapy in Refractory Large B-Cell Lymphoma. New England Journal of Medicine 2017;377(26):2531–2544.

11. Shah NN, Fry TJ. Mechanisms of resistance to CAR T cell therapy. Nat Rev Clin Oncol 2019;16(6):372–385.

12. Donk NWCJ van de, Themeli M, Usmani SZ. Determinants of Response and Mechanisms of Resistance of CAR T-cell Therapy in Multiple Myeloma. Blood Cancer Discov 2021;2(4):302–318.

13. Ruella M, Maus MV. Catch me if you can: Leukemia Escape after CD19-Directed T Cell Immunotherapies. Comput Struct Biotechnol J 2016;14:357–362.

14. Plaks V et al. CD19 target evasion as a mechanism of relapse in large B-cell lymphoma treated with axicabtagene ciloleucel. Blood 2021;138(12):1081–1085.

15. Majzner RG, Mackall CL. Tumor Antigen Escape from CAR T-cell Therapy. Cancer Discov 2018;8(10):1219–1226.

16. Perna F, Sadelain M. Myeloid leukemia switch as immune escape from CD19 chimeric antigen receptor (CAR) therapy [Internet]. Translational Cancer Research 2016;5(2). doi:10.21037/9016

17. Miao L, Zhang Z, Ren Z, Tang F, Li Y. Obstacles and Coping Strategies of CAR-T Cell Immunotherapy in Solid Tumors. Frontiers in Immunology 2021;12:1862.

18. Ambrose C et al. Anti-CD19 CAR T cells potently redirected to kill solid tumor cells. PLoS One 2021;16(3):e0247701.

19. Rennert PD et al. Anti-CD19 CAR T Cells That Secrete a Biparatopic Anti-CLEC12A Bridging Protein Have Potent Activity Against Highly Aggressive Acute Myeloid Leukemia In Vitro and In Vivo. Mol Cancer Ther 2021;20(10):2071–2081.

20. Klesmith JR et al. Retargeting CD19 Chimeric Antigen Receptor T Cells via Engineered CD19-Fusion Proteins. Mol Pharm 2019;16(8):3544–3558.

21. Vosjan MJWD et al. Nanobodies Targeting the Hepatocyte Growth Factor: Potential New Drugs for Molecular Cancer Therapy. Mol Cancer Ther 2012;11(4):1017–1025.

22. Tedder TF, Inaoki M, Sato S. The CD19-CD21 complex regulates signal transduction thresholds governing humoral immunity and autoimmunity. Immunity 1997;6(2):107–118.

23. Hossler P, Khattak SF, Li ZJ. Optimal and consistent protein glycosylation in mammalian cell culture. Glycobiology 2009;19(9):936–949.

24. Flintegaard TV et al. N-Glycosylation Increases the Circulatory Half-Life of Human Growth Hormone. Endocrinology 2010;151(11):5326–5336.

25. Bork K, Horstkorte R, Weidemann W. Increasing the sialylation of therapeutic glycoproteins: the potential of the sialic acid biosynthetic pathway. J Pharm Sci 2009;98(10):3499–3508.

26. Liu X et al. Improved anti-leukemia activities of adoptively transferred T cells expressing bispecific T-cell engager in mice. Blood Cancer J 2016;6(6):e430.

27. Choi BD et al. CAR-T cells secreting BiTEs circumvent antigen escape without detectable toxicity. Nat Biotechnol 2019;37(9):1049–1058.

28. Kim J, et al. Albumin turnover: FcRn-mediated recycling saves as much albumin from degradation as the liver produces. American Journal of Physiology-Gastrointestinal and Liver Physiology 2006;290(2):G352–G360.

29. Ginaldi L et al. Levels of expression of CD19 and CD20 in chronic B cell leukaemias. J Clin Pathol 1998;51(5):364–369.

30. Lemoine J, Ruella M, Houot R. Born to survive: how cancer cells resist CAR T cell therapy. Journal of Hematology & Oncology 2021;14(1):199.

31. Majzner RG et al. Tuning the Antigen Density Requirement for CAR T-cell Activity. Cancer Discov 2020;10(5):702–723.

32. Ramakrishna S et al. Modulation of Target Antigen Density Improves CAR T-cell Functionality and Persistence. Clin Cancer Res 2019;25(17):5329–5341.

33. Pont MJ et al. γ-Secretase inhibition increases efficacy of BCMA-specific chimeric antigen receptor T cells in multiple myeloma. Blood 2019;134(19):1585–1597.

34. Yoshida T et al. All-trans retinoic acid enhances cytotoxic effect of T cells with an anti-CD38 chimeric antigen receptor in acute myeloid leukemia. Clin Transl Immunology 2016;5(12):e116.

35. Walker AJ et al. Tumor Antigen and Receptor Densities Regulate Efficacy of a Chimeric Antigen Receptor Targeting Anaplastic Lymphoma Kinase. Mol Ther 2017;25(9):2189–2201.

36. Watanabe K et al. Target antigen density governs the efficacy of anti-CD20-CD28-CD3 ζ chimeric antigen receptor-modified effector CD8+ T cells. J Immunol 2015;194(3):911–920.

37. Shah N, Chari A, Scott E, Mezzi K, Usmani SZ. B-cell maturation antigen (BCMA) in multiple myeloma: rationale for targeting and current therapeutic approaches. Leukemia 2020;34(4):985–1005.

38. Teoh PJ, Chng WJ. CAR T-cell therapy in multiple myeloma: more room for improvement. Blood Cancer J. 2021;11(4):1–18.

39. Zhang Z et al. Point mutation in CD19 facilitates immune escape of B cell lymphoma from CAR-T cell therapy. J Immunother Cancer 2020;8(2):e001150.

40. Xu X et al. Mechanisms of Relapse After CD19 CAR T-Cell Therapy for Acute Lymphoblastic Leukemia and Its Prevention and Treatment Strategies [Internet]. Frontiers in Immunology 2019;10.https://www.frontiersin.org/article/10.3389/fimmu.2019.02664. cited April 16, 2022

41. Mueller KT et al. Cellular kinetics of CTL019 in relapsed/refractory B-cell acute lymphoblastic leukemia and chronic lymphocytic leukemia. Blood 2017;130(21):2317–2325.

42. Kochenderfer JN et al. Long-Duration Complete Remissions of Diffuse Large B Cell Lymphoma after Anti-CD19 Chimeric Antigen Receptor T Cell Therapy. Mol Ther 2017;25(10):2245–2253.

43. Frigault MJ, Maus MV. State of the art in CAR T cell therapy for CD19^+^ B cell malignancies. J Clin Invest 2020;130(4):1586–1594.

44. Hay KA et al. Factors associated with durable EFS in adult B-cell ALL patients achieving MRD-negative CR after CD19 CAR T-cell therapy. Blood 2019;133(15):1652–1663.

45. Ogasawara K et al. Population Cellular Kinetics of Lisocabtagene Maraleucel, an Autologous CD19-Directed Chimeric Antigen Receptor T-Cell Product, in Patients with Relapsed/Refractory Large B-Cell Lymphoma. Clin Pharmacokinet 2021;60(12):1621–1633.

46. Horna P, Nowakowski G, Endell J, Boxhammer R. Comparative Assessment of Surface CD19 and CD20 Expression on B-Cell Lymphomas from Clinical Biopsies: Implications for Targeted Therapies. Blood [published online ahead of print: 2019]; doi:10.1182/blood-2019-129600

47. Cortelazzo S, Ponzoni M, Ferreri AJM, Hoelzer D. Lymphoblastic lymphoma. Crit Rev Oncol Hematol 2011;79(3):330–343.

48. Frank MJ et al. Monitoring of Circulating Tumor DNA Improves Early Relapse Detection After Axicabtagene Ciloleucel Infusion in Large B-Cell Lymphoma: Results of a Prospective Multi-Institutional Trial. JCO 2021;39(27):3034–3043.

49. Ogawa M, Yokoyama K, Imoto S, Tojo A. Role of Circulating Tumor DNA in Hematological Malignancy. Cancers (Basel) 2021;13(9):2078.

50. Kuhnl A et al. Early FDG-PET response predicts CAR-T failure in large B-cell lymphoma. Blood Adv 2022;6(1):321–326.

51. Milone MC, Bhoj VG. The Pharmacology of T Cell Therapies. Mol Ther Methods Clin Dev 2018;8:210–221.

52. Faude S et al. Absolute lymphocyte count proliferation kinetics after CAR T-cell infusion impact response and relapse. Blood Advances 2021;5(8):2128–2136.

53. Demaret J et al. Monitoring CAR T-cells using flow cytometry. Cytometry Part B: Clinical Cytometry 2021;100(2):218–224.

54. Shalabi H et al. Sequential loss of tumor surface antigens following chimeric antigen receptor T-cell therapies in diffuse large B-cell lymphoma. Haematologica 2018;103(5):e215– e218.

55. Reinhard K et al. An RNA vaccine drives expansion and efficacy of claudin-CAR-T cells against solid tumors. Science 2020;367(6476):446–453.

56. Wang C et al. Anti-PD-1 antibodies as a salvage therapy for patients with diffuse large B cell lymphoma who progressed/relapsed after CART19/20 therapy. Journal of Hematology & Oncology 2021;14(1):106.

57. Chong EA et al. PD-1 blockade modulates chimeric antigen receptor (CAR)–modified T cells: refueling the CAR. Blood 2017;129(8):1039–1041.

58. Rafiq S, Hackett CS, Brentjens RJ. Engineering strategies to overcome the current roadblocks in CAR T cell therapy. Nat Rev Clin Oncol 2020;17(3):147–167.

59. Mehta PH et al. T Cell Fitness and Autologous CAR T Cell Therapy in Haematologic Malignancy [Internet]. Frontiers in Immunology 2021;12.https://www.frontiersin.org/article/10.3389/fimmu.2021.780442. cited April 16, 2022

