## Supplemental Figure 1 for "CAR-T Engager Proteins Optimize Anti-CD19 CAR-T Cell Therapies for Lymphoma"

**Supplemental Figure 1**. Characterization of cell line antigen expression *in vitro* and *in vivo*. A) Expression of the CD19 and CD20 cell surface antigens on Ramos and Jeko-1 cell lines used in the studies. B) Expression of the CD19 and CD20 cell surface antigens on JeKo-19KO cells. C) Growth of different numbers of JeKo-19KO cell implanted sc in NSG mice. Tumors were harvested on day 18. D) Expression of CD19 and CD20 on JeKo-19KO cells after implantation into NSG mice for 18 days compared to wildtype JeKo-1 cells kept in culture (grey lines). E) Dose response curves of the anti-CD20 VHH antibody domain to JeKo-1 and JeKo-19KO cells by flow cytometry.

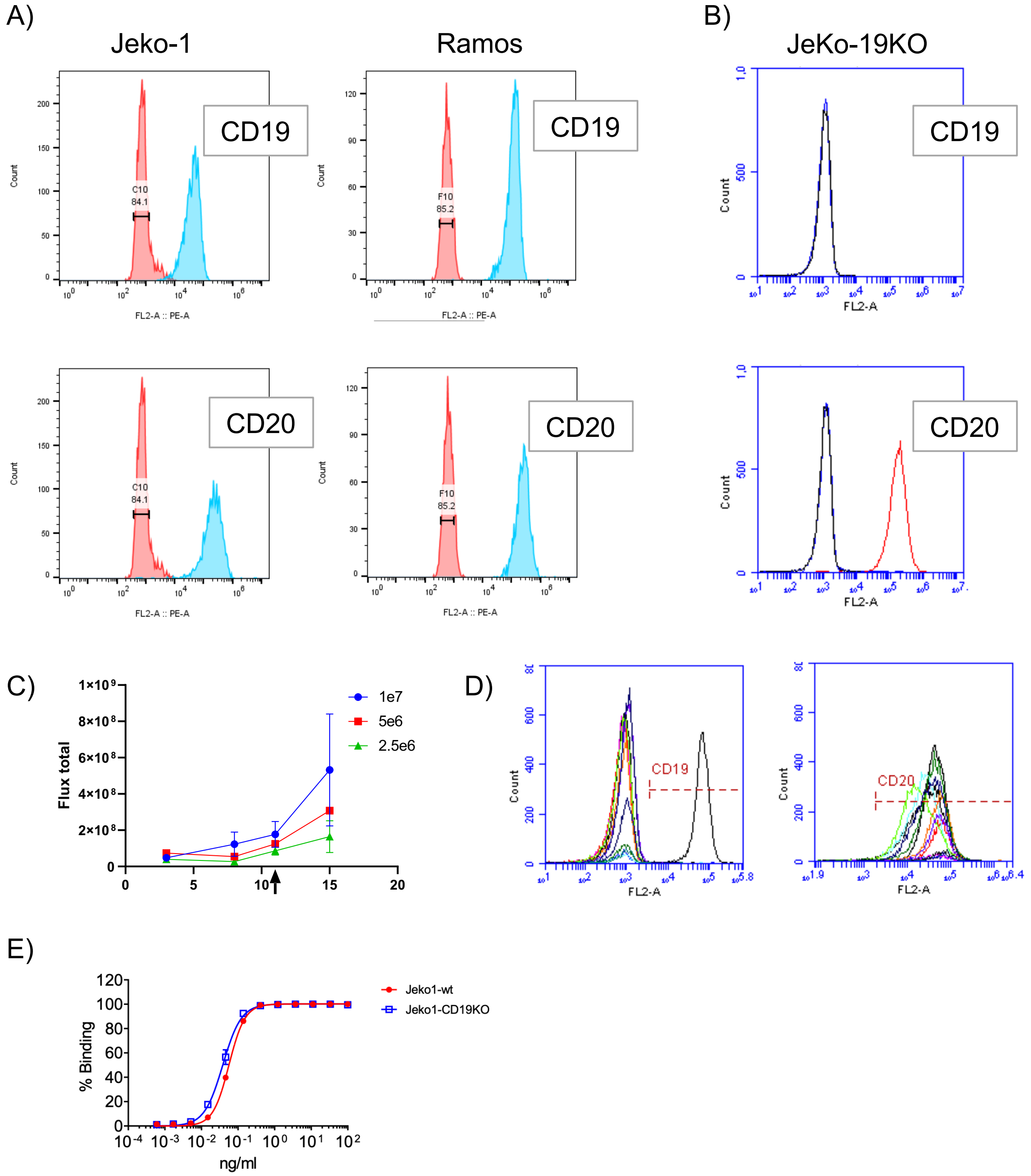
