## Supplemental Figure 2 for "CAR-T Engager Proteins Optimize Anti-CD19 CAR-T Cell Therapies for Lymphoma"

**Supplemental Figure 2**. Protein characterization. A) 4 μg of protein were loaded per lane. CTE-19.20-His (lanes 1-2) and CTE-1.20-RG (lanes 3-4) were run on 4-12% Bis-Tris SDS-PAGE gels. Samples were either non-reduced (lanes 1 and 3) or reduced (lanes 2 and 4). The gels was stained with coomassie blue. B) SEC trace of purified CTE-19.20-His. C) SEC trace of CTE-19.20-RG.

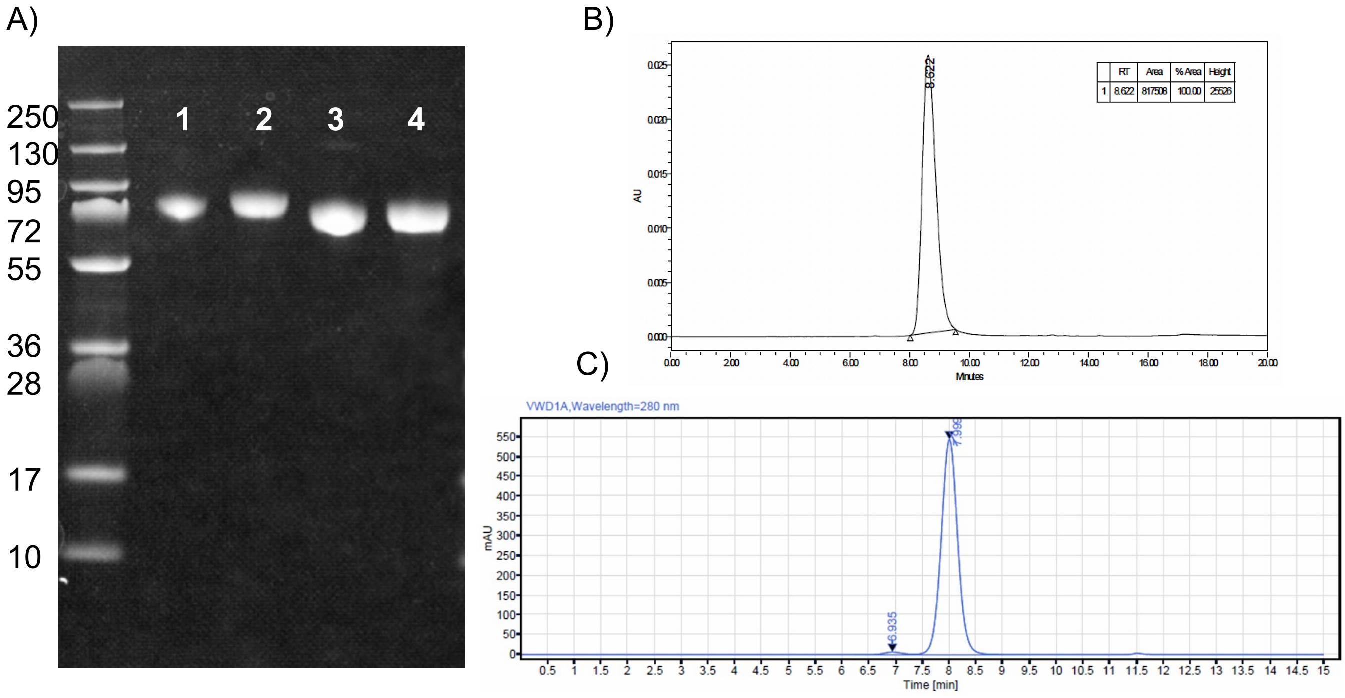
