## Supplemental Figure 3 for "CAR-T Engager Proteins Optimize Anti-CD19 CAR-T Cell Therapies for Lymphoma"

**Supplemental Figure 3**. Excess CTE-19.20-RG protein does not interfere with direct CAR-19 cytotoxicity. CAR-19-mediated killing of JeKo-1 cells was evaluated plus or minus CTE-19.20 protein addition at a) E:T ratio of 3:1 and B) E:T ratio of 1:1. CAR T cells were 78% CAR-positive).

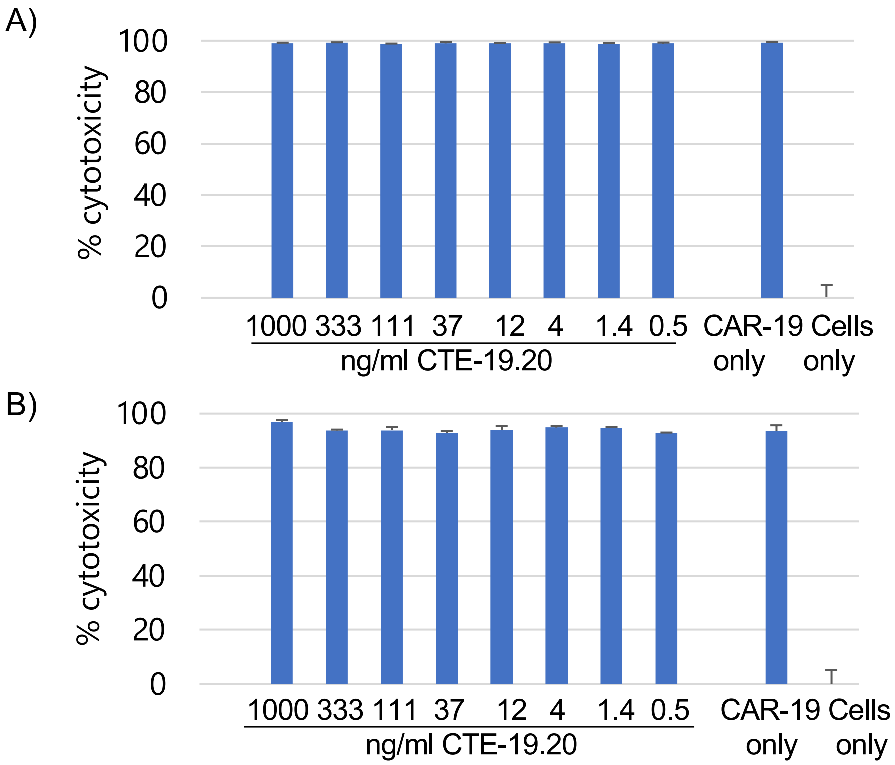
