## Supplemental Figure 4 for "CAR-T Engager Proteins Optimize Anti-CD19 CAR-T Cell Therapies for Lymphoma"

**Supplemental Figure 4.** CTE-19.20 protein binding does not induce CD20 downregulation. A) 1 μg/ml of CTE-19.20-His was incubated for varying amount of time with JeKo-19KO cells at 4$^{o}$C (control) or 37$^{o}$C (test). The cells were washed once and stained with anti-CD19 antibody. B) Mean fluorescence intensity (MFI) of CD19 detection increased during incubation at 37$^{o}$ from 15 minute to 6 hours.

A)

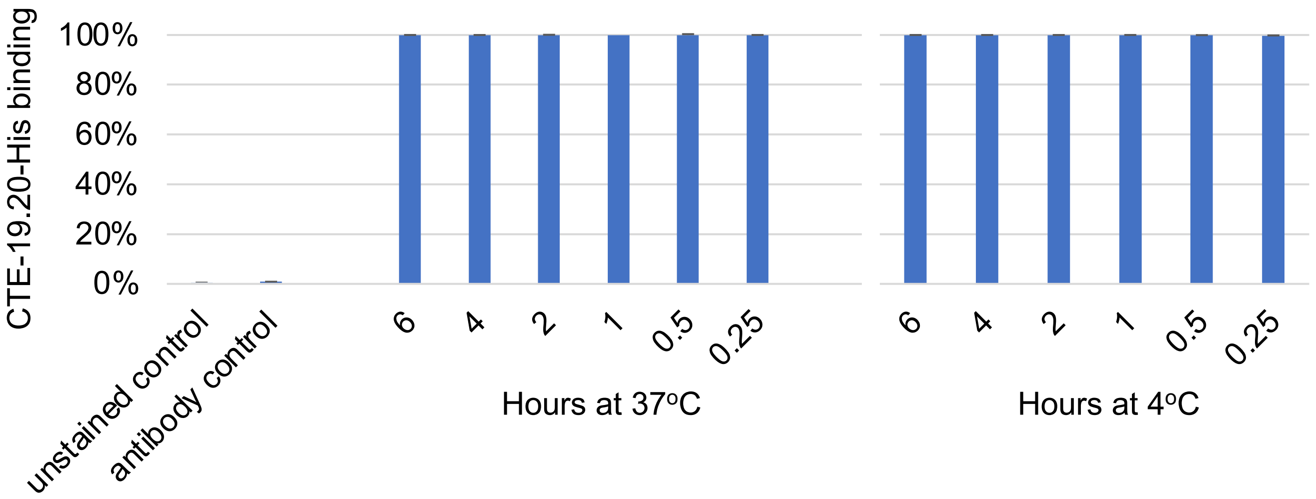


B)

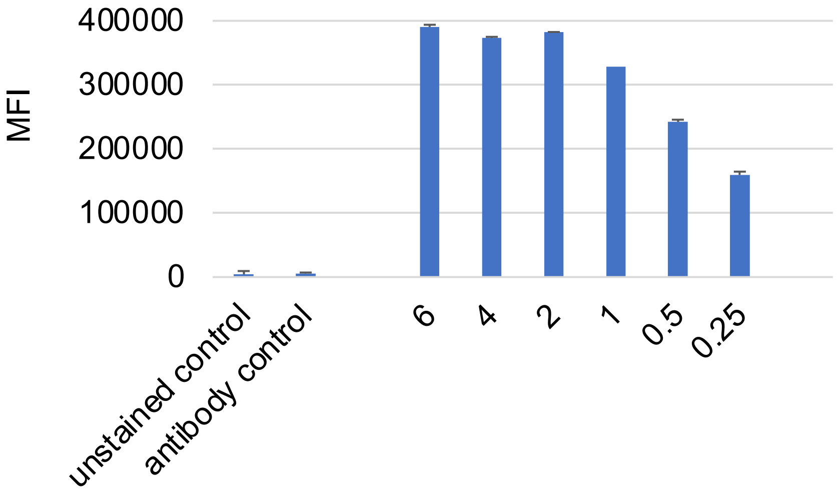
