## Supplemental Figure 5 for "CAR-T Engager Proteins Optimize Anti-CD19 CAR-T Cell Therapies for Lymphoma"

**Supplemental Figure 5.** The cytotoxicity assay was conducted with CAR-19 to target cell ratios of 3:1 (panel A) and 1:1 (panel B). The culture conditions were A) JeKo-19KO + CTE-19.20 + CAR-19 added simultaneously, B) JeKo-19KO + CTE-19.20 preincubated, then added to CAR-19, C) CAR-19 + CTE-19.20 preincubated, then added to JeKo-19KO, D) JeKo-19KO cells alone, E) JeKo-19KO + CAR-19 cells, F) JeKo-19KO + CTE-19.20. * $p < 0.001$ for all comparisons of A, B, or C vs D, E, or F. The CAR-19 T cells were 78% CAR-positive.

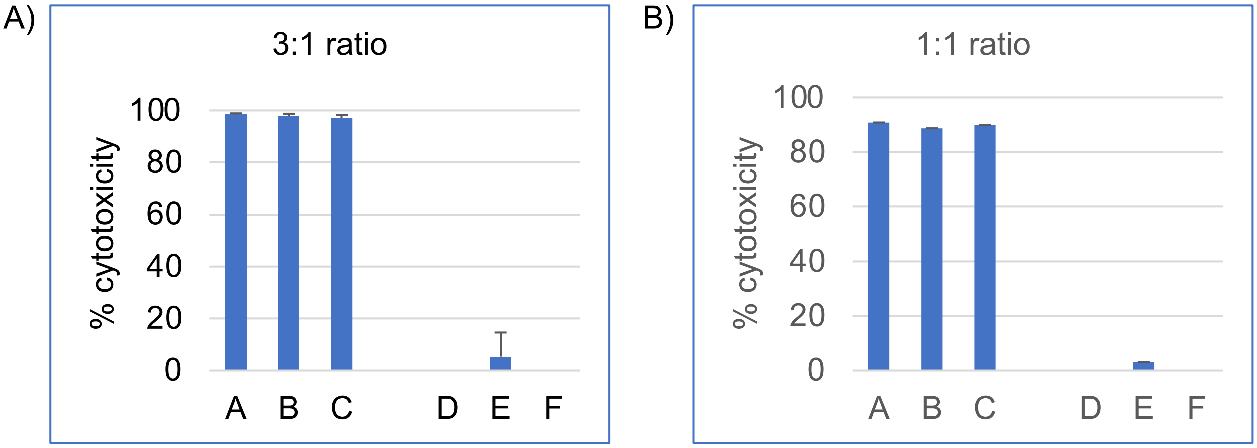
