## Supplemental Figure 6 for "CAR-T Engager Proteins Optimize Anti-CD19 CAR-T Cell Therapies for Lymphoma"

**Supplemental Figure 6**. Binding of CTE-19.20-His to murine and human antigens. A) 293T cells transfected with murine CD20 cDNA express mCD20 as assessed using an anti-mCD20 antibody (left) but do not bind to CTE-19.20 across a range of concentrations from 2 to 0.02 μg/ml. B) The positive control samples using 293 T cells transfected with human CD20 are shown. C) CTE-19.20-His protein binds to murine albumin in an ELISA assay.

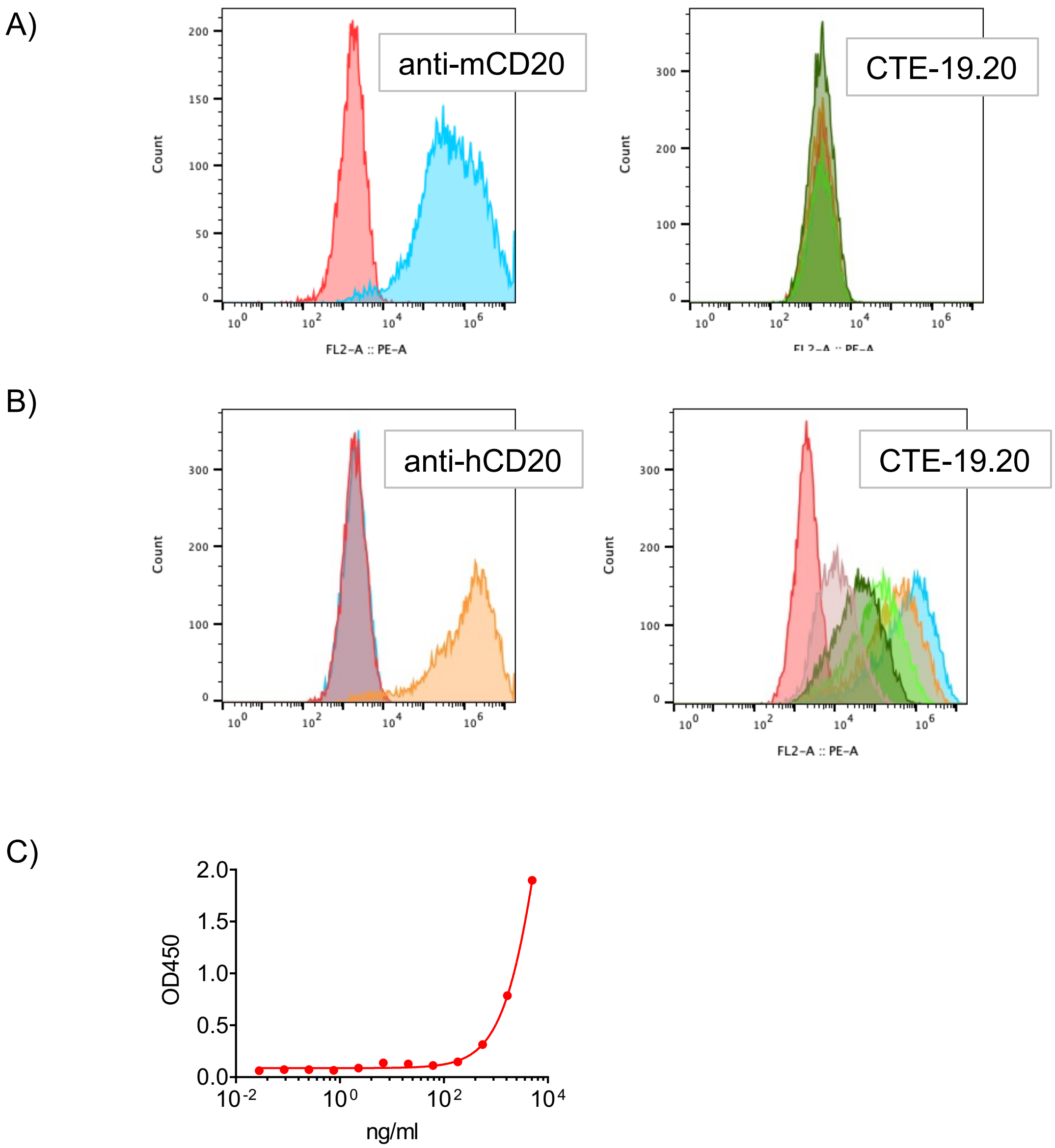
