## Supplemental Table 1 for "CAR-T Engager Proteins Optimize Anti-CD19 CAR-T Cell Therapies for Lymphoma"

**Supplemental Table 1.** Expression of anti-CD19 CAR domains and the CD8 cell surface protein on lentivirus-transduced T cells from two representative donor cell transductions, and cytotoxic activity of the resulting CAR-19 T cells against JeKo-1 cells that express CD19.

| **Donor #** | **Expression** | | **Cytotoxicity** | | |
|---|---|---|---|---|---|
| | % CAR+ | % CD8+ | E:T 3:1 | E:T 1:1 | E:T 0.3:1 |
| **18** | 84.6 | 59.4 | 99.2 | 96.5 | 77.1 |
| **45** | 81.4 | 58.3 | 99.5 | 97.7 | 85.1 |
